# Investigating the Effect of Visual Threat in Virtual Reality on Perceiving Postural Instability Onset

**DOI:** 10.1101/2023.11.17.567574

**Authors:** Robert E McIlroy, Michael Barnett-Cowan

## Abstract

**Background:** Sensory information processing plays a crucial role in monitoring the timing of external and internal events, including the control of balance. While previous research has investigated the role of vision in the perceived timing of postural instability onset with eyes closed and open, it is important to further explore the influence of visual context. Virtual reality offers a unique opportunity to manipulate visual information and assess its impact on balance control and the perceived timing of sensory events.

**Research Question:** Does visual information, particularly visual threat presented in virtual reality, alter the perceived timing of postural instability onset?

**Methods:** Two temporal order judgment tasks were conducted using virtual reality to manipulate visual information. Participants were placed on a virtual skyscraper to induce visual threat. The experiments investigated the impact of visual information on the perceived onset of postural instability while manipulating the presence/absence of visual threat.

**Results:** With vision available but without visual threat, the onset of a postural perturbation needed to occur 10.71-12.33 ms before a reference sound stimulus to be perceived as simultaneous. With visual threat, the onset needed to occur 4.45 ms before auditory cue onset to be perceived as simultaneous. While these delays were not significantly different from true simultaneity of perturbation and sound onset, participants were significantly more precise in their judgments when threatening visual information was present.

**Significance:** Our results show that visual context, particularly visual threat presented in virtual reality, may alter the perception of perturbation onset and the precision of judgments made. This has implications for understanding the role of vision in balance control and developing interventions to improve balance and prevent falls.

## 1. Introduction

Sensory modalities each play a unique role in forming our perception of events and our subsequent actions. Each sensory modality - vision, proprioceptive and vestibular - provides meaningful information to the central nervous system (CNS), facilitating the formation of accurate internal and external models for perception and action. These cues are critical in responding to multisensory events such as postural control. Extensive research has underscored the importance of sensory feedback in controlling posture, highlighting its potential to reduce the risk of falls and injury. Recent studies have also begun to unravel how sensory information may influence our temporal perception during postural tasks, enhancing our understanding of how different types of sensory feedback can impact our perception of postural events. In this context, our study assesses the effect of visual threat on the perception of postural instability onset.

Measuring temporal perception of sensory cues is an effective method for understanding how individuals utilize different sensory modalities to form perceptions of both internal and external events. Temporal order judgment (TOJ) tasks serve as a valuable tool in this regard, offering insights into how individuals perceive information from different sensory modalities. Initial research into the perception of postural events suggested that postural perturbations are perceived more slowly than auditory cues [1,2]. However, more recent studies, taking into account methodological design, have reported no significant effect of visual information on the perceived timing of postural perturbations, challenging previous findings [3,4]. Nevertheless, other research in postural control has shown that visual context, such as height, can substantially influence an individual’s perception and actions [5-18]. Assessing the impact of altering visual context on the perceived timing of perturbation onset, therefore, could provide valuable insights into how threatening visual information is integrated into balance control.

Visual information plays a critical role in providing visuospatial cues about our external environment, cues that can subsequently influence motor commands related to postural control. Studies have shown that visuospatial information provided prior to a postural perturbation can directly affect change-in-support postural strategies, such as reach-to-grasp [19] and stepping [20]. Additionally, altering visual information to create a perceived threat, such as height, can impact postural responses during both quiet stance [5-7] and postural perturbations [8-11], whether in real-world or virtual environments. Visual height has also been observed to influence perceptual measures such as fear, anxiety, and postural confidence [7,12]. These findings highlight the significant role that visual information can play in shaping both postural responses and our perception of postural control. However, it is crucial to note that these research paradigms provided context-specific visual information, whether from objects in the external environment or perceived limitations imposed by the visual environment, such as height. The differentiation of visual context is the primary focus of this research.

This study puts forth three main hypotheses. First, we aim to replicate previous findings that the perception of postural instability onset is not significantly slower than that of an auditory reference stimulus [3,4]. Second, we hypothesize that presenting a visually threatening scene via virtual reality (VR) will reduce any perceived delay between postural instability onset and an auditory reference stimulus compared to a non-threatening environment, based on observed behavioral and psychological changes at both physical and virtual heights [7,11-18]. Finally, we anticipate that participants will exhibit improved precision or a reduction in the variability of their responses, resulting in a decreased just noticeable difference (JND) in their TOJ tasks when exposed to a virtual height. This hypothesis is grounded in the idea that perceived threat from the VR scene will increase attention, which has been shown to reduce JND in TOJ tasks [21].

## 2.0 Methods

### 2.1 Participants

A total of 14 participants (6 male and 8 female) were recruited, all free of musculoskeletal, auditory, visual, vestibular, or other neurological disorders. One participant was removed from the analysis because they were not able to complete all trials in the experiment, resulting in a final sample of 13 participants (6 male and 7 female; aged 19-24). All participants gave their informed written consent in accordance with the guidelines set by the University of Waterloo Research Ethics Committee.

### 2.2 Protocol

The apparatus, stimuli and procedure used were consistent with previous work conducted within our lab [3,4], with the exception of utilizing a virtual reality head mounted display (VR HMD) for one of the conditions.

This study’s methodology mirrored that of previous research conducted within our lab [3,4]. Participants made temporal order judgment (TOJ) responses to a postural perturbation evoked by a lean-and-release mechanism and an auditory reference cue. The setup for the lean-and-release and auditory reference stimulus was identical to previous research [3,4]. Participants indicated their responses regarding which stimulus occurred first using two handheld buttons. The two conditions in this study were eyes open (EO) and virtual reality (VR). During EO trials (non-threatening visual context), participants were instructed to focus on an ‘x’ marked on a wall approximately 3 meters away within the lab space [22]. During VR trials (threatening visual context), participants wore a VR headset and were positioned at the edge of a tall virtual building in a city setting (Figure 1). They were instructed to fix their gaze on the top of the virtual building in front of them to limit eye movement during the trials. The stimulus onset asynchrony (SOA) distribution used in this experiment was designed to be equal (cf [3,4]). Participants completed 144 trials (72 trials with eyes open and 72 trials with VR), with each SOA repeated eight times. The SOAs ranged from -200 ms to 200 ms, the first increment was 25 ms, with each subsequent SOA doubling in size resulting in steps at 0 ms, ±25ms, ±50 ms, ±100 ms and ±200 ms. [3,4]. Negative SOAs indicate that the postural perturbation occurred first, and positive SOAs indicate that the auditory reference stimulus occurred first. The SOAs were fully randomized for each participant. After every 24 consecutive trials, participants were given a 3-minute break to sit down, preventing fatigue over the course of the study. Each trial lasted 15-20 seconds, with a comparable delay between each trial. Participants were instructed to relax, keeping their arms and hands comfortably at their sides throughout the trials. Prior to the experiment, participants completed five practice trials.

**Figure 1:**
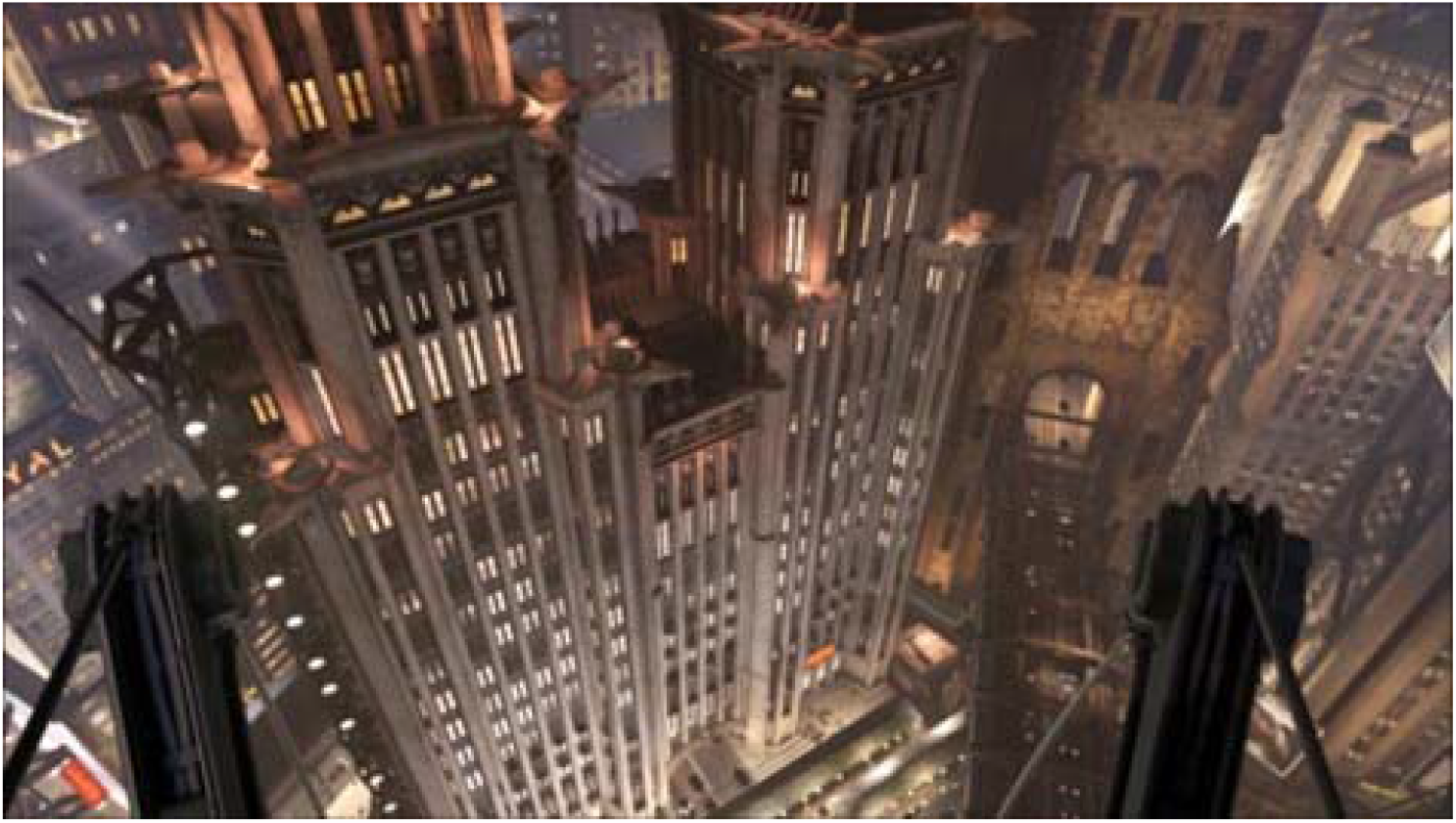
Virtual scene presented to the participants during the VR condition. Participants were placed at the edge of a virtual skyscraper while physically leaning forward with the lean-and-release paradigm.

Participants’ TOJ responses were to a postural perturbation evoked by a lean-and-release mechanism and an auditory reference stimulus delivered through headphones. Participants were first weighed to calculate 7-8% of their body mass. This value was then used to calculate the corresponding force that would equate to this 7-8% body mass. Participants then adopted a lean angle that matched this force value on the load cell. They were fitted with a full-body harness allowing for the attachment of the lean cable at the level of the 2nd and 3rd thoracic vertebrae and the safety rope, which was secured to the ceiling to prevent injury in case of an inability to recover balance [1,2,22,23]. Participants were positioned approximately 1 m from the lean-and-release apparatus, with their feet in a standardized position (heel centers 0.17 m apart, 14° between the long axes of feet [1,2,24]. The ground was marked with tape along the lateral borders of the feet, and a piece of wood served as a heel stop to ensure the foot position remained constant during the study. Participants received no specific instructions on how to respond to the postural perturbation, to prevent any voluntary changes to their postural response. The lean angle of 7-8% body mass produced a perturbation large enough to inherently evoke a stepping response in each subject, but it was not small enough to permit a fixed support postural strategy [1,2,23,25,26].

### 2.3 Virtual Reality Head Mounted Display and Visual Scene

This experiment used an Oculus Rift CV1 VR HMD during VR conditions. The headset displayed a resolution of 960 X 1080 pixels to each eye, resulting in a total screen resolution of 1920 × 1080 pixels. The system had a refresh rate of 90 Hz and a 100-degree horizontal field of view. The lenses were adjusted to fit each participant’s eyes, and the support straps were adjusted for a comfortable fit.

### 2.4 Lean-and-Release Apparatus & Stimuli

The lean-and-release mechanism and the auditory stimuli were identical to those used in previous research; please refer to this prior work for specific characteristics of the system [3,4].

### 2.5 Data Analysis

The data analysis procedure was identical to that performed in previous research, including the fitting of the logistic function (Eq. 1) [3,4]. The same criteria were applied to determine if a participant’s data would be included in our final analysis [3,4].

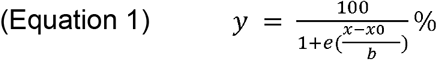

### 2.6 Statistical Analysis

A within-subjects design was used to reassess the hypothesis that postural instability does not need to occur prior to an auditory reference stimulus for it to be perceived as simultaneous. A one-sample t-test was conducted to compare the mean point of subjective simultaneity (PSS) value of each condition relative to true simultaneity (0 ms). Paired sample t-tests were performed across the EO and VR conditions to determine if a virtual threat caused a decrease in both PSS and just noticeable difference (JND) means. Additionally, an independent paired-samples t-test was performed to compare the PSS and JND means of the EO condition from previous research using the same paradigm and methodology with the EO condition from this study. This comparison aimed to determine whether different populations of participants produce consistent results for the same perceptual task. If normality failed in any comparisons, as assessed by the Shapiro-Wilk test, we attempted to normalize the data. If Z-Score normalization was unsuccessful, non-parametric statistical tests were performed. For each of the statistical tests, a significance level of α = 0.05 was used.

## 3. Results

During our analysis, we excluded one participant’s data as it did not satisfy the criteria related to the logistic regression fit [3,4]. Specifically, the fit of the VR data points yielded an R^2^ = 0.42 and a p-value = 0.0585, indicating a poor fit and unreliable parameter estimates.

Figure 2 displays the logistic fits for each participant (grey lines), as well as the averaged logistic fit across all participants (black line), for each condition. We derived the PSS and JND values for all participants from these fits (see Figure 2). For the EO condition, the results were a mean PSS of -10.71 ms (SE = 11.02, median PSS = -10.47 ms), while for the VR condition, the mean PSS was -4.45 ms (SE = 6.44, median PSS = -9.98 ms). Both conditions produced average PSS values near true simultaneity (Figure 3), indicating that in the EO condition, the postural instability onset needed to precede the auditory cue by 10.71 ms to be perceived as simultaneous. In the VR condition, this requirement was 4.45 ms. The VR PSS mean was closer to true simultaneity, showing a slight reduction compared to the EO PSS.

**Figure 2:**
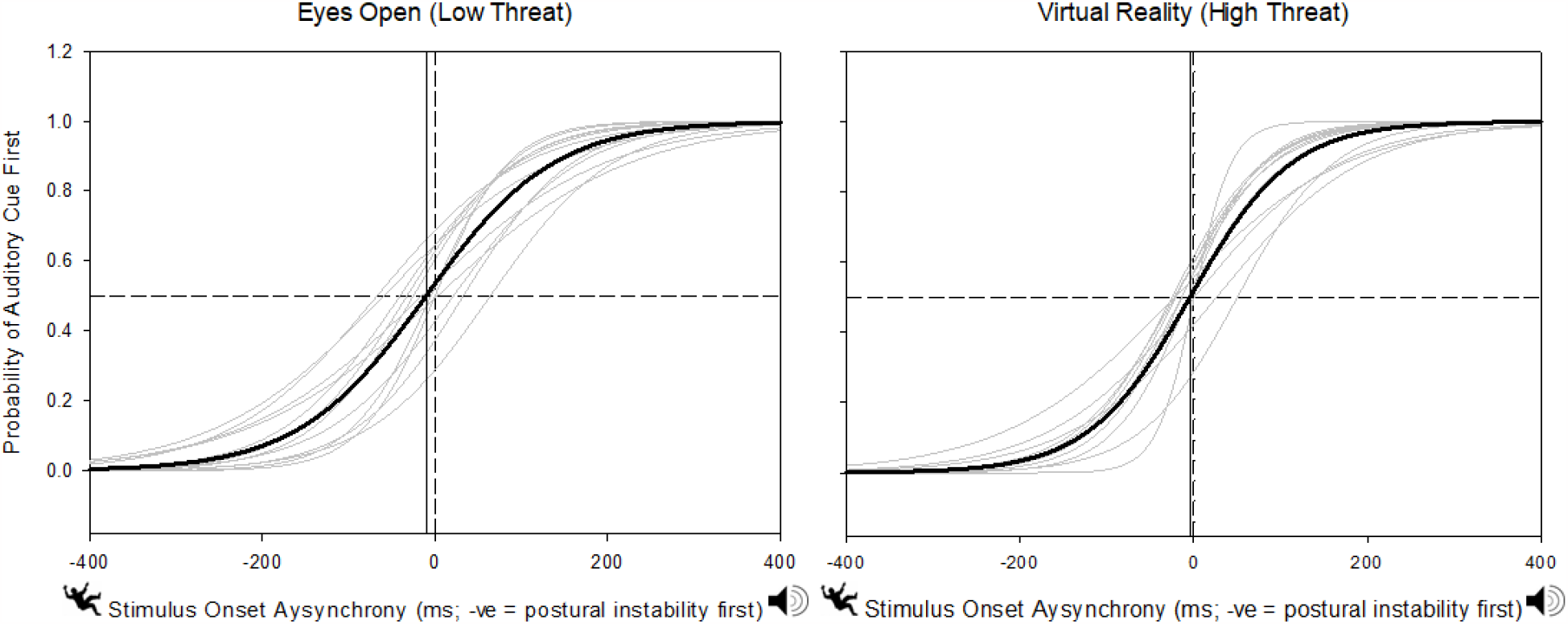
Logistic fits for the Eyes Open versus Virtual Reality conditions. Light grey lines represent individual logistic function fits, while the black line represents the mean. The solid vertical line shows the mean PSS value for each condition. A 0 ms SOA, marked by a vertical dashed line, denotes true simultaneity between stimuli, and the horizontal dotted line represents the 50% probability of the auditory reference stimulus occurring first, corresponding to the PSS.

**Figure 3:**
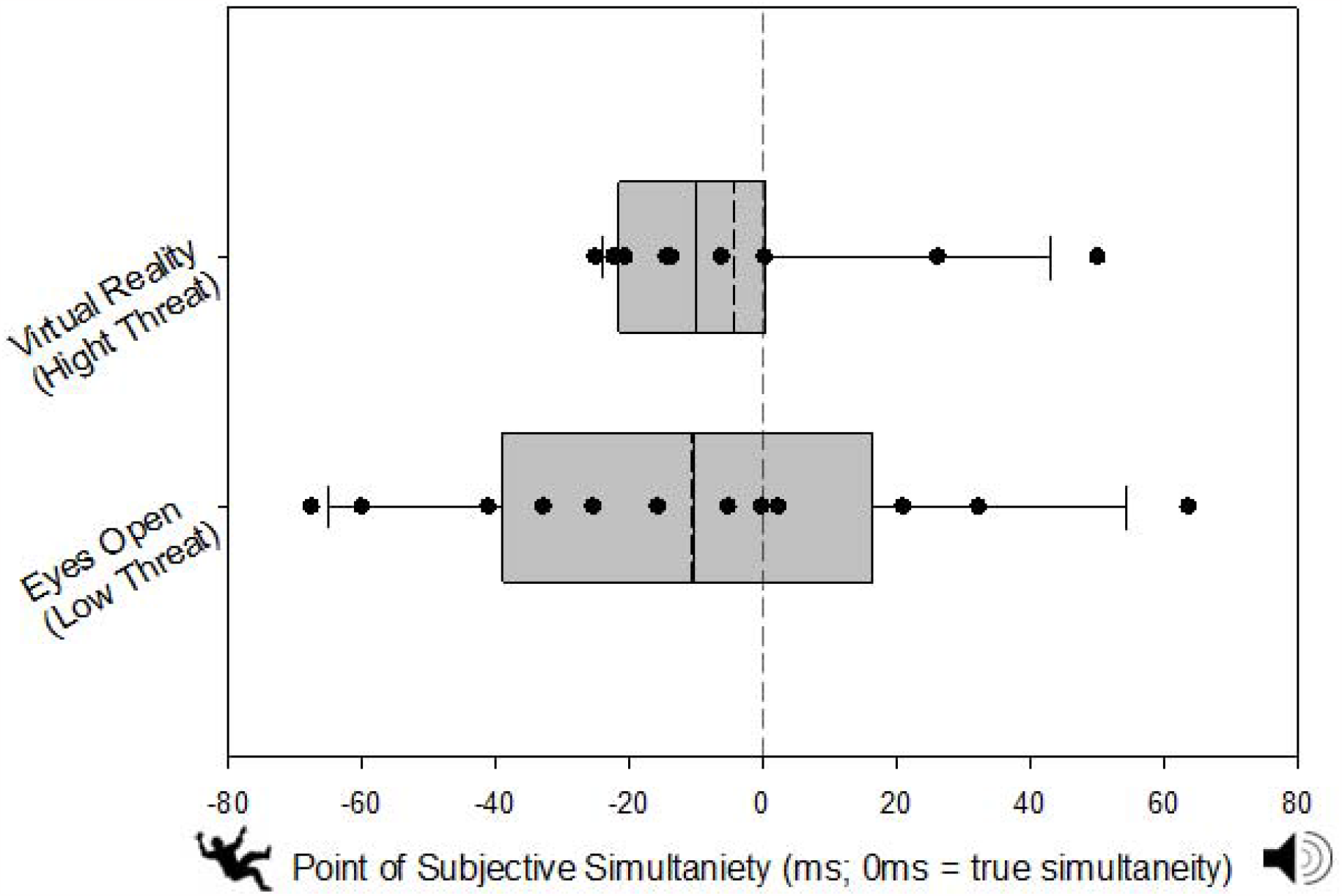
Mean PSS (dashed line), median PSS (solid line), and individual PSS values (circles) for both the VR (top) and EO (bottom) conditions. Grey bars indicate the 25-75th percentiles, and error bars show the 10th and 90th percentiles of the PSS data. The VR and EO conditions’ PSS values showed no statistically significant differences.

One-sample t-tests on the EO PSS (t(11) = -0.972; p = 0.176) and the VR PSS (Z = -1.020; p = 0.170) showed that the PSS from each condition was not significantly different from 0 ms, aligning with previous findings where PSS means did not significantly deviate from true simultaneity [3,4]. Additionally, a paired sample t-test between the EO and VR PSS means (t(11) = -0.798; p = 0.442) revealed no statistically significant difference, suggesting that the VR’s threatening visual context did not notably affect the PSS means compared to the non-threatening condition.

As shown in Figure 4, the EO and VR conditions’ mean JND values (EO JND = 73.99 ms, SE = 6.41, median = 69.83 ms; VR JND = 58.31 ms, SE = 5.95, median = 53.73 ms) differed by 15.68 ms, a significant change of 26.89% (t(11) = 3.416; p = 0.006; 1 - β = 0.64). This finding implies that a threatening VR environment resulted in participants responding more precisely and consistently.

**Figure 4:**
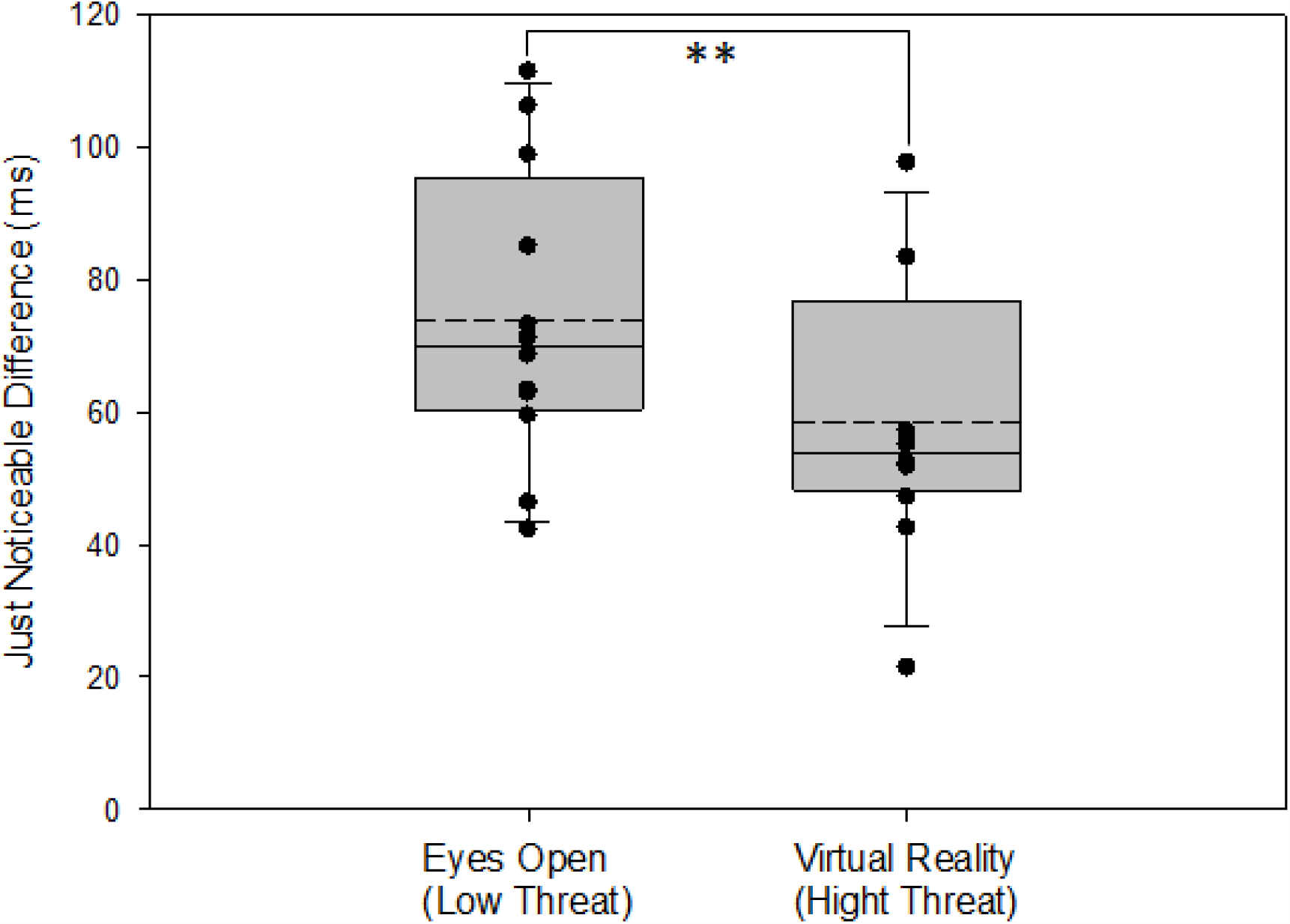
Mean JND (dashed line), median JND (solid line), and individual JND values (circles) for the EO (left) and VR (right) conditions. Grey bars indicate the 25-75th percentiles, and error bars show the 10th and 90th percentiles of the JND data. The VR and EO conditions’ JND means differed significantly, with a p-value < 0.01 indicated by **.

## Discussion

This study investigated the impact of visual context, particularly threatening visual stimuli presented in VR, on the perceived simultaneity between postural instability onset and an auditory reference stimulus. As hypothesized and corroborated by previous research [3,4], visual information—whether obtained simply by opening the eyes or within VR—did not significantly alter the PSS relative to true simultaneity (0ms). Surprisingly, introducing a virtual threatening scene in VR also did not significantly affect the PSS, contradicting our initial hypothesis, which expected the threat to reduce perceived delay of perturbation onset.

However, the JND values in the VR condition showed a substantial 26.89% decrease compared to the eyes open condition, confirming our hypothesis and indicating enhanced precision of responses when exposed to the virtual threatening scene.

These consistent findings, demonstrating no perceptual delay in perceiving the onset of postural instability, validate the use of equal SOA distribution to prevent lag adaptation from asynchronous stimuli presentation [3,4,27-31]. This underscores the significance of our chosen methodology in perceptual research and attests to its consistency and reproducibility across different participant groups.

Concerning visual context, introducing a threatening virtual height seemed to necessitate a heightened awareness for detecting perturbation onset during the VR condition, compared to a normal, eyes-open state in the lab. Yet, this did not translate to significant differences in the PSS means, suggesting that perception of postural instability onset may predominantly rely on proprioceptive feedback. This is likely due to proprioception’s rapid processing speeds [32,33], its substantial role in balance control [34], and its precision in detecting self-motion [35].

On the other hand, the JND (precision) of responses markedly differed between the eyes-open and VR conditions. Research has consistently shown that VR and threatening visual scenes can induce psychological effects such as increased arousal, heightened anxiety, and diminished confidence in balance [7,11-18]. These effects likely sharpen an individual’s attention, which, while not affecting PSS, has been found to improve the precision (reduce JND) of TOJ responses [21]. Li et al. (2018) attributed this to phasic arousal—arousal shifts induced by stimulus conditions. Moreover, emotionally evocative images have been found to enhance temporal precision [36], suggesting a top-down influence of evocative visual cues on cortical regions involved in TOJ task precision.

Studies employing functional magnetic resonance imaging and magnetoencephalography have explored the cortical substrates of various sensory TOJ tasks, identifying involvement of areas such as the temporal parietal junction [37], supramarginal gyrus [37,38], right posterior parietal cortex [39], bilateral inferior parietal cortices [38], superior and middle temporal gyrus [38], insula [37,38], and frontal gyri [37,38]. This body of research points to the inferior parietal lobe as a likely site for TOJ decision-making. Traditionally, visual information is processed through dorsal and ventral pathways—the former dealing with spatial perception and motion in parietal regions, and the latter with object recognition in temporal regions [40]. However, Battelli et al. (2007) proposed a ‘when’ visual pathway related to timing or perception of events, localized to the right inferior parietal lobe—coincidentally, the same region activated during TOJ tasks. This suggests that visual information could modulate this cortical region during TOJ tasks, potentially explaining the increased precision observed in the VR condition.

A limitation of this study is that the non-threatening visual stimuli were not presented in VR. Consequently, sensory differences due to wearing the VR headset, such as its weight, the vividness of imagery, or the sense of presence in the virtual environment, should be taken into account. These factors could be assessed using questionnaires during and after data collection or controlled for by using VR scenes in both conditions.

In conclusion, while visual context does not seem to affect perceived onset of postural instability, it significantly influences response precision. These findings, consistent with results obtained with other types of visual stimuli, pave the way for future research to explore perceptual responses in older adults and assess the potential of this task for fall risk quantification. Further investigations could also examine how visual context and other sensory modifications influence perception of postural instability in specific populations.

## Conflict of Interest

There are no conflicts of interest.

## Acknowledgements

Supported by a Natural Sciences and Engineering Research Council of Canada (NSERC) Discovery Grant (RGPIN-03977-2020) and an Ontario Early Researcher Award to MB-C.

## Notes

### Competing Interest Statement

The authors have declared no competing interest.

